# Schizophrenia Risk Mapping and Functional Engineering of the 3D Genome in Three Neuronal Subtypes

**DOI:** 10.1101/2023.07.17.549339

**Authors:** Samuel K. Powell, Will Liao, Callan O’Shea, Sarah Kammourh, Sadaf Ghorbani, Raymond Rigat, Rahat Elahi, PJ Michael Deans, Derek J. Le, Poonam Agarwal, Wei Qiang Seow, Kevin C. Wang, Schahram Akbarian, Kristen J. Brennand

## Abstract

Common variants associated with schizophrenia are concentrated in non-coding regulatory sequences, but their precise target genes are context-dependent and impacted by cell-type-specific three-dimensional spatial chromatin organization. Here, we map long-range chromosomal conformations in isogenic human dopaminergic, GABAergic, and glutamatergic neurons to track developmentally programmed shifts in the regulatory activity of schizophrenia risk loci. Massive repressive compartmentalization, concomitant with the emergence of hundreds of neuron-specific multi-valent chromatin architectural stripes, occurs during neuronal differentiation, with genes interconnected to genetic risk loci through these long-range chromatin structures differing in their biological roles from genes more proximal to sequences conferring heritable risk. Chemically induced CRISPR-guided chromosomal loop-engineering for the proximal risk gene *SNAP91* and distal risk gene *BHLHE22* profoundly alters synaptic development and functional activity. Our findings highlight the large-scale cell-type-specific reorganization of chromosomal conformations at schizophrenia risk loci during neurodevelopment and establish a causal link between risk-associated gene-regulatory loop structures and neuronal function.

## INTRODUCTION

Schizophrenia risk is highly heritable^1^, with a complex polygenic architecture reflecting the contributions of dozens of rare coding variants that confer relatively large impacts^2^ together with hundreds of non-coding common variants that contribute individually small effect sizes^3^. Common risk variants are enriched in regions that regulate gene expression during neurodevelopment and in the adult brain^4–7^, particularly within highly organized three-dimensional (3D) genome structures^8^. We previously monitored developmentally regulated changes in neuronal and glial 3D chromatin structures, demonstrating neural cell-type-specific coordination at the level of the chromosomal connectome, transcriptome, and proteome^9^. Chromosome conformations and chromatin structures vary considerably by cell type^10^, even among specific subtypes of neurons^11^, but to date, these structures have been studied with genome-wide coverage in relatively few subpopulations of the human brain^8, 12–18^. Although global chromosome conformations are critically important for brain development and function^19^, their individual causal roles in neurodevelopment and disease remain unresolved.

Here we constructed 3D genome maps of isogenic induced dopaminergic (iDOPA), GABAergic (iGABA), and glutamatergic (iGLUT) neurons generated from human induced pluripotent stem cells (hiPSCs). The hiPSC-to-neuron transition was marked by substantial genome-wide expansion of repressive (“B”) chromatin compartments; unexpectedly, a subset of persistently active (“A”) compartments were enriched in schizophrenia risk loci and harbored neuron-specific chromatin architectural stripes (“stripes”) that only emerged upon differentiation. Stripes occur when specific loop anchors interact across an entire region at high frequency^20, 21^, facilitating transcription through an association to active enhancers and super-enhancers, and are largely under explored in the brain^22–24^. Here, neuronal stripes were enriched at sites harboring brain-specific super-enhancers, with loops connecting schizophrenia risk loci to distal neurodevelopmentally regulated genes.

We identified a disproportionate number of neuronal subtype-specific stripes and loops, which were significantly enriched for genes associated with chromatin regulation, cell adhesion, and synaptic functions. Site-specific, dCas9-mediated experimental induction of schizophrenia-associated loops at the proximal risk gene *SNAP91* and distal risk gene *BHLHE22* altered target gene expression and disrupted the structure and function of neuronal networks. Overall, we demonstrate the functional significance and causal impact of cell-type-specific chromatin dynamics across the three neuronal subtypes critically relevant to the pathophysiology of schizophrenia.

## RESULTS

### Large-scale repressive compartmentalization during hiPSC-to-neuron transition spares schizophrenia risk loci

Transient overexpression of *ASCL1, LMX1B,* and *NR4A2* (also known as “Nurr1”) induced iDOPA neurons within 21 days that were 92% positive for TH, synthesized dopamine, were enriched in fetal midbrain dopaminergic neuron gene expression signatures and exhibited electrophysiologic hallmarks of midbrain dopaminergic neurons by day 35^25^. Likewise, transient overexpression of *ASCL1* and *DLX2* induced iGABA neurons within 35 days that were 95-99% positive for GABA itself and GAD1/2, the majority of which (>75%) were of the SST+ subtype and possessed mature physiologic properties of inhibitory neurons by day 42^26–28^. Finally, we^28–32^ and others^33–40^ demonstrated that transient overexpression of *NGN2* induced iGLUT neurons within 21 days that were >95% pure glutamatergic neurons, robustly expressed glutamatergic genes, released glutamate, produced spontaneous synaptic activity, and recapitulated the impact of psychiatric disease associated genes.

The 3D genomes of hiPSCs and hiPSCs-derived iDOPA, iGABA, and iGLUT neurons were segmented into active (“A”) compartments and inactive (“B”) compartments using principal component analysis (PCA) of Hi-C eigenvector scores from two independent donors. Each sample showed distinct clusters by cell type identity; hiPSCs separated clearly from the three neuronal subtypes on principal component 1 (“PC1”) explaining 33.6% of the variance, and specific neuronal subtype clusters separated on principal component 2 (“PC2”) accounting for 22.4% of the variance (**Figure 1A**). We assessed the similarity between samples, at the donor and cell type level, using stratum adjusted correlation coefficient (SCC) metrics^41^ (**Supplementary Figure 1**). There was substantially higher SCC within cell types compared to within donors (*p* = 1.6×10^-4^), as well as between same cell-types compared to different cell types. Given this and the observed clustering of samples in PCA by cell type independent of donors, we combined data from different donors for each cell type to create high coverage Hi-C maps and downsampled them each to 185 *cis* interactions for further analysis (**Supplementary Dataset 1B**).

**Figure 1.**
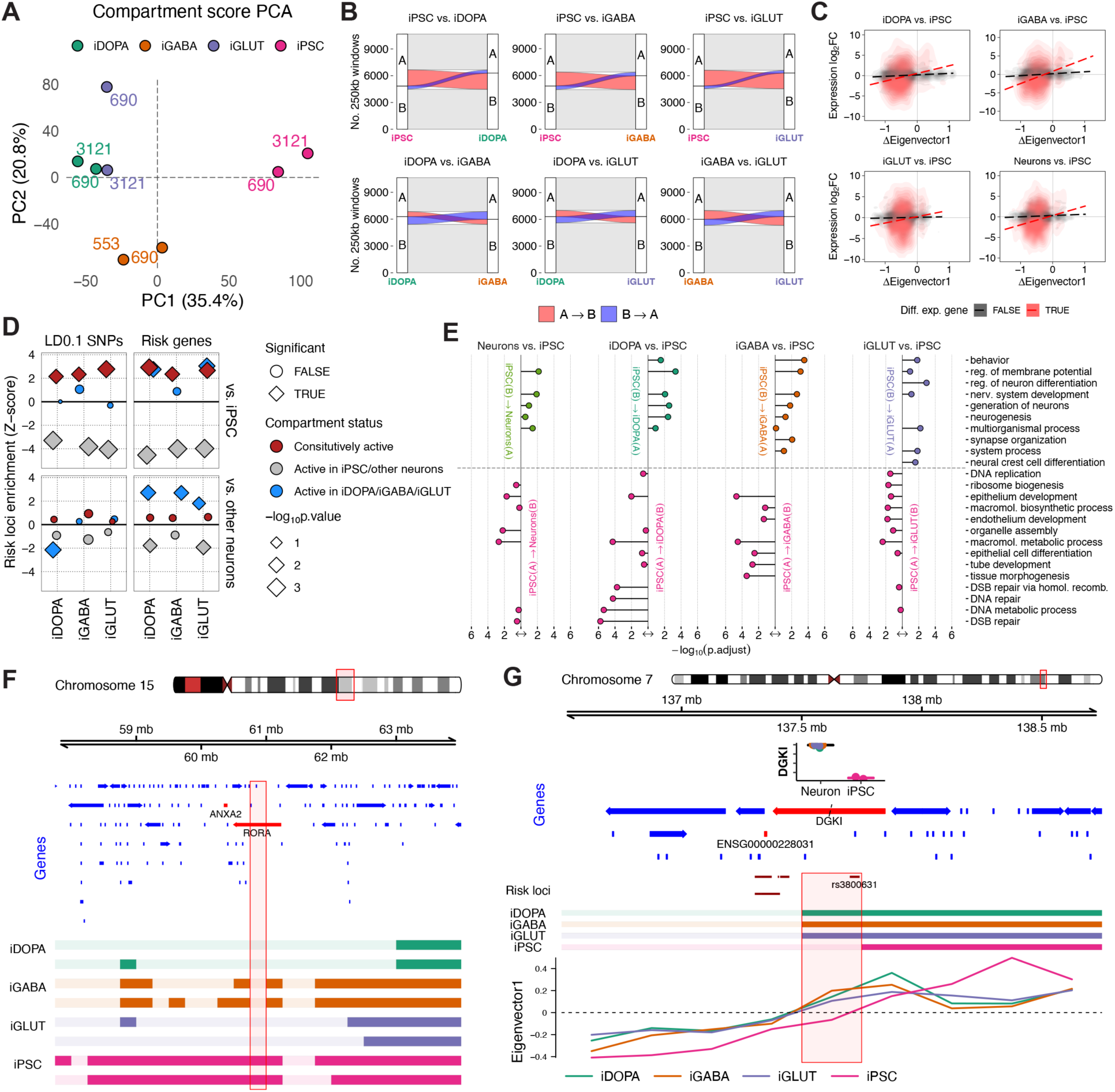
Chromosomal Compartment Conversion During Neurodevelopment Correlates with Neuronal Subtype-Specific Gene Expression and Spares SCZ Risk Loci. **(A)** Genome-wide eigenvector score for N=8 Hi-C samples from control donors 553, 690, and 3121 as indicated, showing clustering by cell type. **(B)** Genome-wide A/B compartment numbers and proportions, (top) hiPSC-to-iDOPA, -iGABA and -GLUT differentiation as indicated, (bottom) comparison between neuronal subtypes. Notice massive A -> B flux during hiPSC-to-neuron transition. **(C)** Cell-specific differential gene expression correlates with score differential of Hi-C eigenvector. **(D)** Schizophrenia PGC3 risk locus enrichment (Z) by cell and compartment type, notice significant risk locus depletion in differentiation-induced B-compartment sequences. **(E)** Gene Ontologies for 250kb compartment blocks undergoing A -> B or B -> A switching during hiPSC-to-neuron differentiation, as indicated. **(F,G)** Compartment status switch from A in hiPSCs to B in three neuronal subtypes at the (F) *RORA* and (G) *DGK1 risk loci*.

Compartment switching (**Supplementary Dataset 2**), assessed at 250kb resolution in the merged Hi-C maps during hiPSC-to-neuron differentiation revealed large-scale B-compartmentalization (389-454Mb, or 12.6-14.7% of total genome sequence, dependent on neuronal subtype). In striking contrast (*p* = 4.9×10^-324^, binomial test), only a very minor portion (2.5-3.4% of the genome) converted to active A-compartment status during hiPSC-to-neuron differentiation (**Figure 1B**). In general, developmentally regulated compartment switching was similar across iDOPA, iGABA, and iGLUT neurons, with limited A/B compartment differences between the three neuronal subtypes (A-to-B 4.6-8.0% and B-to-A 5.5-7.0% of genome) (**Figure 1B**). Chromosomal compartment architectures within a given cell type were highly reproducible across donors, the fraction of concordant compartment calls between same cell types was significantly greater then between different cell types (*p* = 9.4×10^-3^) (**Supplementary Figure 2)**, and as illustrated by a representative region on chromosome 15 around the nuclear hormone receptor and neurodevelopmental risk gene, *RORA* (**Figure 1F**). Dynamic changes in A/B compartmentalization during hiPSC-to-neuron differentiation were broadly correlated to transcriptomic changes (**Supplementary Dataset S5**) of the corresponding genes (**Figure 1C**), with functional enrichment for neuron-specific gene ontologies and de-enrichment for genes regulating tissue morphogenesis and early development (**Figure 1E**).

The developmentally regulated A/B compartment map was super-imposed with the most recent genome wide association study (GWAS) map for schizophrenia^3^, comprised of 291 common risk loci and 1111 PsychENCODE schizophrenia risk genes. Across all three neuronal subtypes, schizophrenia-associated single nucleotide polymorphisms (SNPs) (linkage disequilibrium (LD)>0.1, n = 139167) and gene target(s) were depleted in regions undergoing B-compartmentalization during hiPSC-to-neuron differentiation (compared to randomly sampled, GC-content matched windows, n = 1000, *p* = 1.0×10^-3^ for each neuronal subtype) (**Figure 1D**). Instead, schizophrenia risk was significantly enriched in neuronal A compartment chromatin (*p* = 3.0×10^-3^ – 2.0×10^-2^), as is the case for the chr7 rs3800631 risk SNP within the *DGKI* gene (**Figure 1G**)).

Taken together, while neuronal induction of hiPSCs was marked by substantial genome-wide expansion of repressive (“B”) compartments, sequences that maintained active (“A”) compartment status during hiPSC-to-neuron differentiation were significantly enriched for schizophrenia risk loci.

### Risk-associated stripes overlap with brain-specific super-enhancers at sites of developmentally regulated gene expression

In peripheral cells, stripes represent sequentially ordered contacts generated at domain boundaries anchored at cohesion docking sites, mark regulatory hubs such as super-enhancers^23^, and drive coordinated expression of developmentally programmed gene expression and cell-type identity^42^. We detected stripes on our high coverge, merged data using Stripenn^43^ and identified 217 iDOPA, 575 iGABA, 193 iGLUT, and 623 hiPSC stripes^43^. Stripe coordinates were then consolidated by merging overlapping stripes into a single, non-contiguous set (1,071 stripes spanning 717Mb). Finally, the contact enrichment (observed/expected, O/E, contact frequencies) of each donor-level Hi-C dataset was scored over the new coordinates (**Supplementary Dataset 3A**). PCA was then performed on donor-level O/E frequencies across each of the stripes for each sample (**Figure 2B**) which revealed the expected separation of neurons from hiPSCs samples. We observed a prominent example of a stripe, called in the merged neuron Hi-C contact matrix, with broad enrichment between the *PTPRU* risk gene and a nearby brain super-enhancer on chromosome 1 (**Figure 2A).**

**Figure 2.**
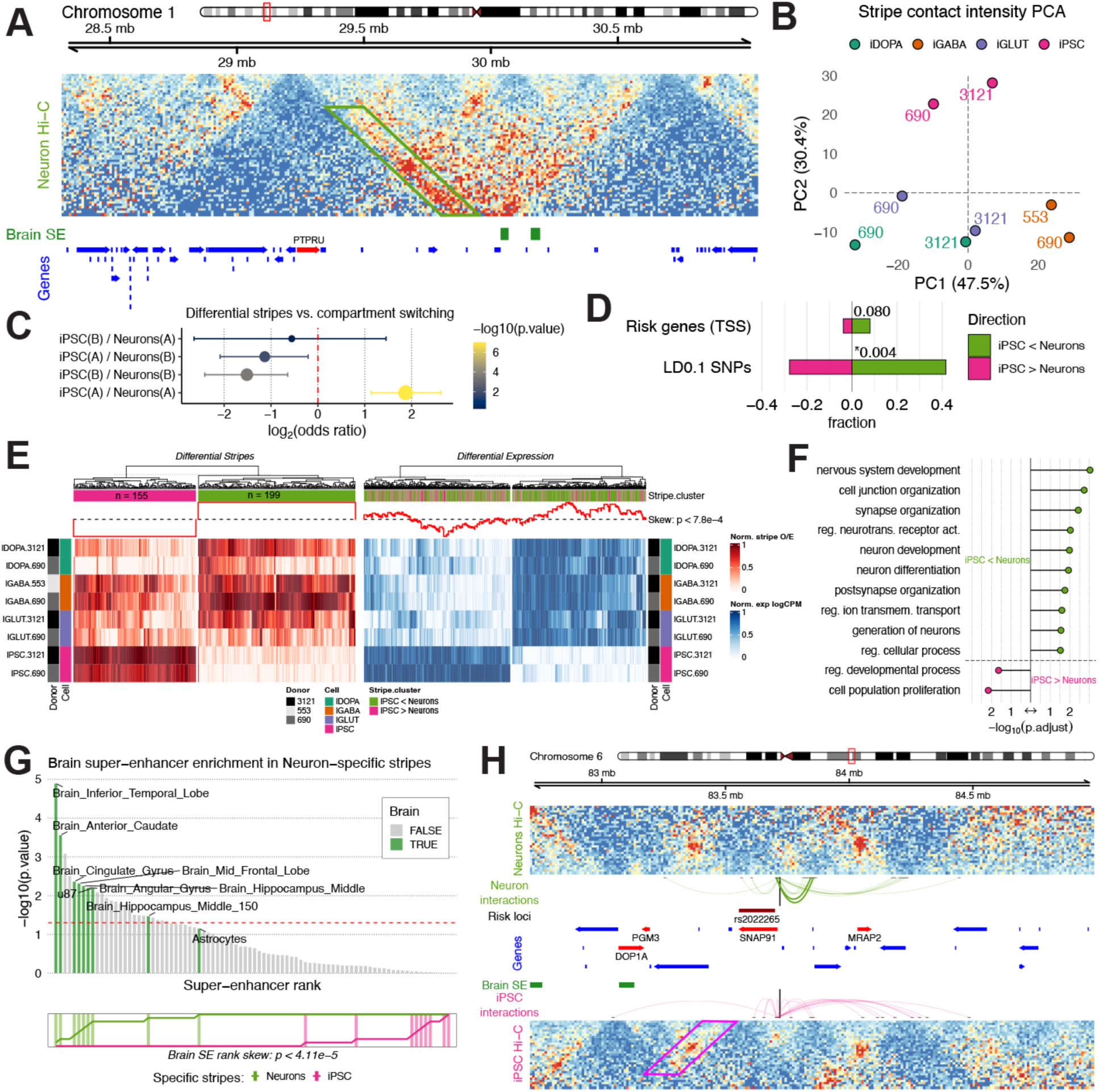
Chromatin Stripes Structurally Connect Neurodevelopmentally Regulated Gene Expression and Schizophrenia Risk Loci. **(A)** Example neuron-specific stripe anchored at the schizophrenia risk gene *PTPRU*. **(B)** PCA of Hi-C samples show clustering by cell type. **(C)** Neuron-specific stripes are enriched in chromatin compartments that are active (A) in hiPSCs and remain active upon neuronal differentiation. **(D**) Enrichment of risk loci among neuron-specific stripes. **(E)** Hierarchical clustering of Hi-C samples by hiPSC-specific versus neuron-specific stripes and strong correlation between stripe specificity and differential gene expression. **(F)** Genes contained within neuron-specific stripes are enriched in biological pathways regulating neurodevelopment and synapse organization. **(G)** Neuron-specific stripes are enriched in brain tissue super enhancer regions. **(H)** Emergence of neuronal subtype-specific loops targeting schizophrenia risk gene *SNAP91* neighboring neuronal stripes.

We identified cell-specific stripes by performing t-tests on donor-level O/E contact frequencies between cell types (**Supplementary Dataset 3B**) and found that risk loci falling within loci maintaining A compartment status during hiPSC-to-neuron transition frequently were significantly enriched in neuron-specific chromatin architectural stripes (“stripes”) (*p* = 9.5×10^-8^) **(Figure 2C**) and showed significant overlap with PGC3 schizophrenia SNPs (LD>0.1, *p* = 4.3×10^-3^) (**Figure 2D**). Conversely, hiPSC-specific stripes were enriched in neuron-repressed compartments (hiPSC(A)/Neurons(B): *p* = 1.0×10^-2^; hiPSC(B)/Neurons(B): *p* = 2.5×10^-4^) (**Figure 2C**). Neuron-specific stripes were also enriched for super-enhancers (using super-enhancer brain reference maps^44, 45^). Brain-related super-enhancers tended to be significantly more enriched in neuron-specific stripes compared to hiPSC-specific stripes, where they were among the least enriched (*p* = 4×10^-5^, Wilcoxon rank sum test) (**Figure 2G, Supplementary Dataset 3E**). To corroborate the regulatory role of neuron-specific stripes, we examined the abundance of differentially expressed genes (DEGs) in comparison to stripe O/E values. Expectedly, neuron- and hiPSC-specific stripes clustered separately and showed distinct O/E intensities in neuronal or hiPSC Hi-C data. Interestingly, after clustering DEGs based on expression levels across eight RNA-seq samples from the same cell types, genes upregulated in neuronal subtypes were significantly enriched within neuron-specific stripes (*p* = 7.8×10^-4^, Fisher’s test) (**Figure 2E, Supplementary Dataset 3C**); likewise, genes upregulated in hiPSCs were more often associated with hiPSC-specific stripes (*p* = 7.8×10^-4^). The DEGs in neuron-specific stripes were overrepresented for neuron-related gene sets, whereas those in hiPSC-specific stripes were not (**Figure 2F, Supplementary Dataset 3D**).

Taken together, neuron-specific stripes and their loop anchors are regulatory domains for developmentally relevant gene expression at sites of brain-specific super-enhancers and confer heritability risk for schizophrenia.

### Neuron-specific 3D chromatin loops are anchored in schizophrenia risk loci

Loops, defined as point-to-point chromosomal contacts, were called by Peakachu^46^ at 10kb resolution (**Supplementary Dataset 4A**). Consistent with prior observations^47^, hiPSC-specific loop numbers (N = 22211) by far outnumbered their neuronal counterparts (**Figure 3A**). PCA performed on the donor-level O/E contact frequencies at called loops demonstrated clustering broadly by cell type (**Figure 3B**). The presence or absence of a loop in the various cell-type contexts was used to define cell-specific loops. Given the similarities between stripe and loop detection, neuronal sub-type specific loops that were absent in hiPSC were unsurprisingly enriched within subtype-specific stripes (iDOPA-*p* = 3.4×10^-62^, iGABA-*p* = 1.5×10^-26^; iGLUT-specific loops *p* = 4.2×10^-48^) (**Figure 3C**). To assess a potential functional role for promoter-associated loops, we evaluated whether loops contacting differentially expressed gene promoters were more likely to link to enhancers or schizophrenia risk SNPs (LD<0.01) or were more likely to be risk genes (**Figure 3D**). We found specific loops contacted both DEGs and fetal enhancers more than expected in iDOPA (*p* = 2.5×10^-9^), iGABA (*p* = 5.3×10^-9^), and Neurons (*p* = 1.8×10^-7^). Additionally, iGABA loops were significantly enriched for risk genes (*p* = 5.0-x10^-2^).

**Figure 3.**
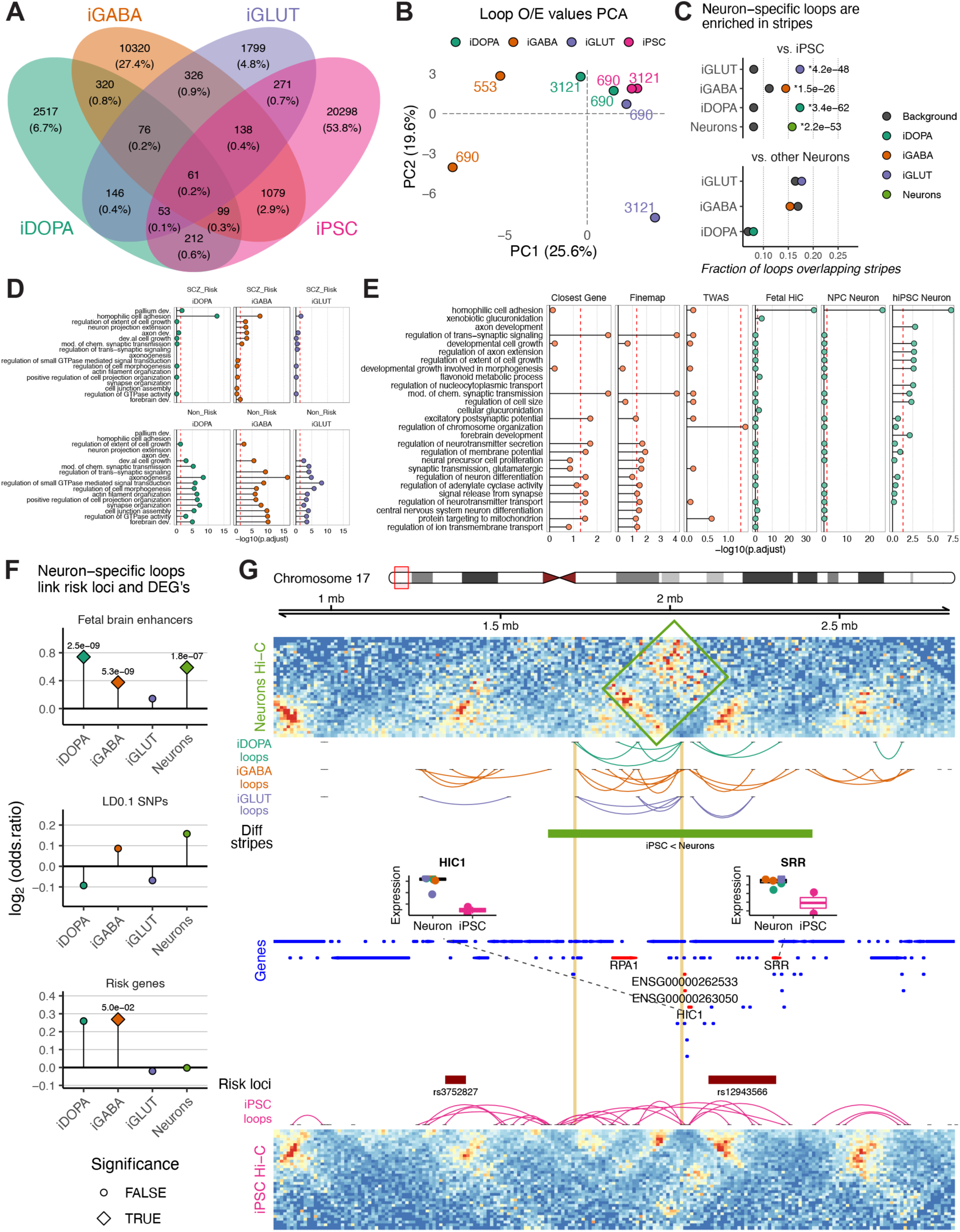
Neuronal Subtype-Specific Loops Are Enriched in Risk Loci Targeting Unique Molecular Pathways. **(A)** Venn diagram showing the number of loops specific to or shared among the different cell types. **(B)** PCA of observed versus expected (O/E) values of loop calls showing clustering of samples by cell type. **(C)** Loops specific to one neuron type or another are enriched among neuronal stripes. **(D)** Genes anchored in chromatin loops differ in biological pathway enrichments based upon risk versus non-risk anchors across neuronal subtypes. **(E)** Distinct pathway enrichments among genes tagged based upon proximity to schizophrenia risk loci versus through long-range chromatin looping. **(F)** Neuronal subtype-specific chromatin loops are enriched in fetal brain enhancers, risk loci, and previously established risk genes. **(G)** Screenshot showing neuron-specific stripe near risk loci with high neuronal express of the genes *HIC1* and *SRR*.

Two broad sets of loops with importance to genetic risk for schizophrenia were identified: the first was defined by an anchor positioned at a gene within a risk locus (“Risk Gene-Connecting Loops”), and the second by a loop connecting a distal gene to a risk locus (“Risk Locus-Connected Genes”) (see Rajarajan et al^9^). This expanded the list of potential risk genes by considering not just linear sequence proximity but also neuronal cell-type-specific 3D spatial proximity, connecting loci as far as 3Mb apart in linear space. On a genome-wide scale, hundreds of Risk Gene-Connecting Loops targeted established schizophrenia risk genes in one or more neuronal subtypes. Neuron subtype-specific (vs. other subtypes) Risk Locus-Connected Genes (N=2201) were specifically enriched in the cell adhesion biological processes (**Figure 3E**) and were more highly enriched for fetal brain-specific enhancers (**Figure 3F**). A representative example is provided in **Figure 3G**, showing the *SRR* serine racemase risk locus.

Taken together, schizophrenia risk loci are enriched for neuronal subtype-specific loops, particularly those linked to enhancers, genes that become activated during neuronal differentiation, and those enriched in cell adhesion.

### Differential enrichment of proximal and distal target genes of schizophrenia risk loci

Linear and 3D chromatin spatial proximity approaches for ascertaining schizophrenia risk genes were contrasted (**Supplementary Dataset 4**). Linearly (ie. proximal) schizophrenia gene targets were: 1) located within PGC3 risk loci (if there were none, we selected the closest gene within 500kb^3^), 2) identified by using Finemap^48^ based on GWAS SNP and brain eQTL-colocalization, and/or 3) predicted using transcriptome-wide association imputation (TWAS) of schizophrenia-associated SNPs from reference post-mortem transcriptomic datasets^49^. In contrast, 3D-defined (ie. distal) schizophrenia risk gene targets were identified by spatial Hi-C chromatin interactions from: 1) fetal cortical plate and germinal zone tissue^8^, 2) NPC-derived *NGN2* neurons^13^, or 3) iDOPA-, iGABA-, and iGLUT-specific loops.

Overall, there were substantial differences in biological processes enriched among proximal and distal gene targets identified **(Figure 3E)**. Proximal risk genes were enriched in brain-related biological processes such as axonogenesis and forebrain development; in contrast, distal risk genes were instead enriched exclusively in homophilic cell adhesion, largely driven by genes from the *protocadherin* gene family.

### Chromatin loop engineering validated functional impact of iGABA-loop targeting SNAP91

To probe schizophrenia risk genes for 3D proximity to distal noncoding regions with high sensitivity, we employed a previously used binomial testing approach to identify interactions with significantly more counts than expected at that distance^9^. We noticed one such interaction, which we interpreted as a loop, connecting the promoter region of the schizophrenia risk gene *SNAP91* to a distal non-coding region approximately 150kb further upstream, which was specific to iGABA neurons. Of note, *SNAP91* is a well-established eQTL-based risk gene for schizophrenia^50^ responsible for the reuptake of vesicle-mediated neurotransmitter and recycling of presynaptic terminals^51–54^. Suppression of *SNAP91* reduces synaptic activity in both animal models^54^ and hiPSC-derived neurons^32^. Therefore, we wanted to recapitulate this iGABA-specific loop by creating it *de novo* in iGLUT neurons, using dCas9 effectors fused to an abscisic acid (ABA)-inducible dimerization system^55, 56^ (**Figure 4A, B, Supplementary Figure 3A-J**).

**Figure 4:**
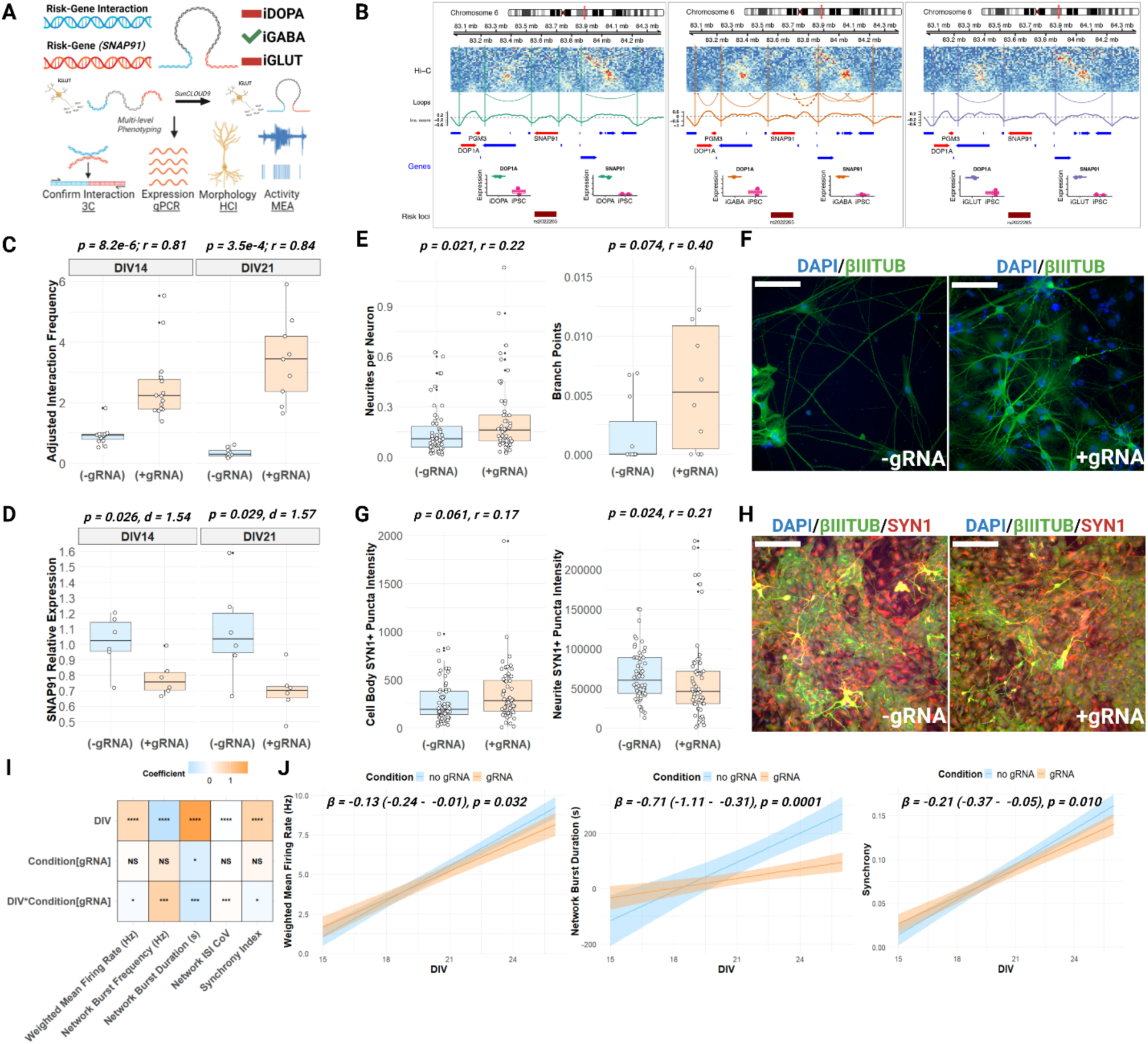
Chromosomal Loop Engineering with Multi-Level Phenotypic Impact at *SNAP91* Risk Gene. **(A)** Schematic for loop induction of an iGABA neuron-specific loop connecting risk-gene-interaction with risk-gene (*SNAP91*) in iGLUT neurons. **(B)** Hi-C maps of the loop involving a distal region and the promoter region of *SNAP91*. **(C)** 3C-PCR demonstrated increased contact frequency between the two bins at DIV14 and DIV21 (n = 2 biological replicates/time point/condition). **(D)** Loop-formation reduced *SNAP91* expression by 25% (normalized to *B-ACTIN* mRNA expression as a loading control) in DIV14 and DIV21 in iGLUT neurons (at least n = 6 replicate wells per timepoint, per DIV, in each condition). **(E-F)** At DIV14, loop-formation increased the number of neurites per neuron and branch points per neurite (n = 4 or more replicate wells with 3 fields per well for each timepoint per condition). **(F)** Representative confocal images of DIV14 SunCLOUD9 iGLUT neurons, ßIII-TUBULIN (green), and DAPI-stained nuclei (blue), scale bar = 50μm. **(G)** Formation of the *SNAP91* loop increased puncta intensity on neuronal cell bodies but decreased puncta intensity on neurites (n = 4 or more replicate wells with 3 fields per well for each timepoint per condition). **(H)** Representative confocal images of DIV21 SunCLOUD9 iGLUT neurons, SYN1 (red), ßIII-TUBULIN (green) and DAPI-stained nuclei (blue), scale bar = 50μm. **(I)** Model coefficients from longitudinal assessment of neuronal activity with and without formation of the *SNAP91* chromatin loop (n = 24 replicate wells per timepoint in each condition). Loop-formation altered maturation-dependent changes in weighted mean firing rate (WMFR), field-level network burst frequency (Hz), network burst duration (s), network inter-spike-interval coefficients of variation (“Network ISI CoV”), and network synchrony. *, p< 0.05, **, p< 0.01, *** p< 0.001. **(J)** Loop-formation at *SNAP91* slowed the maturation-dependent increase in weighted mean firing rate (WMFR) (β = -0.13 (-0.24 - -0.01), p = 0.032), network burst duration (β = -0.71 (-1.11 - -0.31), p = 0.001), and synchrony (β = -0.21 (-0.37 - -0.05), p = 0.010).

**Figure 5.**
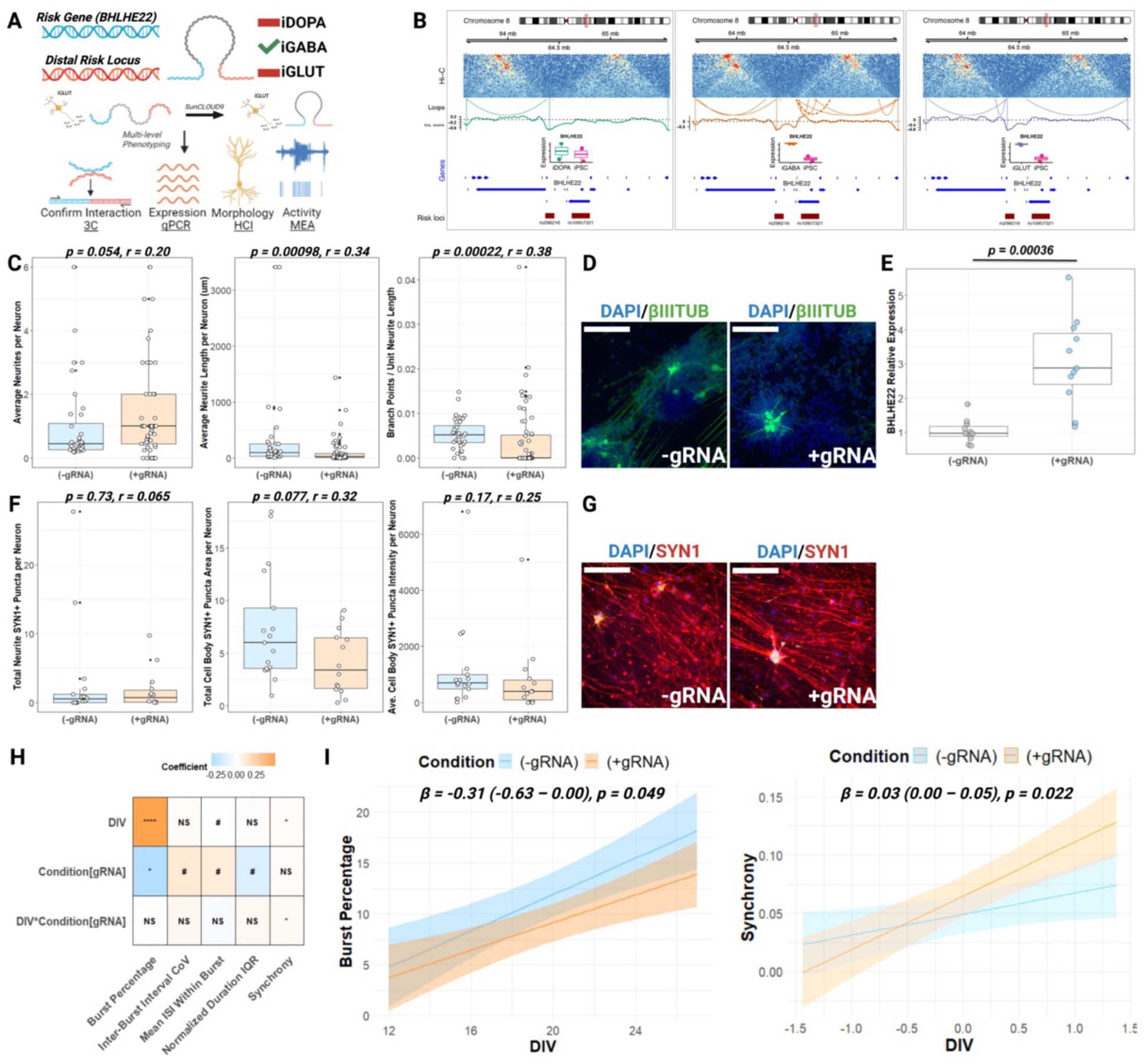
Establishing Enhancer Activity of Risk Locus Targeting *BHLHE22* Through Chromatin Engineering. **(A)** Schematic for induction of an iGABA neuron-specific loop targeting the brain-enriched transcription factor *BHLHE22* through distal looping interactions. **(B)** Hi-C maps of the long-range loop of a risk locus-anchored region targeting the brain-specific transcription factor *BHLHE22* in iGABA neurons only; *BHLHE22* is specifically expressed at high levels in iGABA neurons compared to the other neuron types. **(C)** Loop-formation in iGLUTs increased the average number of neurites per neuron (*p* = 0.054, *r* = 0.20), decreased average neurite length (*p* = 0.0010, *r* = 0.34) and number of branch points (*p* = 0.00013, *r* = 0.39) (n = 4 or more replicate wells with 3 fields per well for each timepoint per condition). **(D)** Representative confocal images of DIV14 SunCLOUD9 iGLUT neurons, ßIII-TUBULIN (green) and DAPI-stained nuclei (blue), scale bar = 50μm. (**E)** Loop creation in iGLUT neurons normally lacking the loop leads to a substantial increase in *BHLHE22* expression by 300%, (normalized to *B-ACTIN* mRNA expression as a loading control), directly demonstrating enhancer activity of the risk locus (n = at least 6 replicate wells per condition). **(F)** No significant change in synaptic puncta number and distribution following loop formation, with representative images shown in **(G). (H)** Summary of model coefficients from longitudinal neuronal activity profiling. (n = 24 wells per condition per timepoint). Loop-formation at *BHLHE22* in SunCLOUD9 iGLUT neurons decreased the percentage of electrodes with bursting activity bursting over neurodevelopment (“burst percentage) without any changes in the frequency (p = 0.34) or duration (p = 0.56) of bursts. There were insignificant differences across measures such as average interval between spikes within bursts (“mean ISI within burst”) (p = 0.077), variability in the time between bursts (“normalized duration IQR”) (p = 0.105), and interburst interval coefficient of variation (“IBI CoV”) (p = 0.057). There was a significant interaction between DIV and treatment condition such that those SunCLOUD9-iGLUTs with gRNAs developed increased synchrony at a faster rate. *, p< 0.05, **, p< 0.01, *** p< 0.001. **(I)** Loop-formation at *BHLHE22* in SunCLOUD9 iGLUT neurons decreased the percentage of electrodes with bursting activity bursting over neurodevelopment (“burst percentage) (β = -0.31 (-0.63 – 0.00), p = 0.049) and increased synchrony at a faster rate (β = 0.03 (0.00 – 0.05), p = 0.022).

At DIV14, SunCLOUD9-iGLUT neurons treated with ABA and transfected with gRNAs had substantially increased interaction frequency between the two 10kb bins constituting the loop compared to a no-gRNA negative control group (Wilcoxon-Mann-Whitney test: *p* = 8.2 x 10^-6^; *r* = 0.814); this effect was maintained at DIV21 (Wilcoxon-Mann-Whitney test: *p* = 3.5 x 10^-4^; *r* = 0.843) (**Figure 4C**). Upon loop-induction, *SNAP91* mRNA expression was decreased by ∼25% at both DIV14 (*p* = 0.026) and DIV21 (*p* = 0.029) compared to SunCLOUD9-iGLUTs receiving a no-gRNA empty transfection **(Figure 4D**). There was no evidence that docking of the SunCLOUD9 effectors to the two loci impacted *SNAP91* expression compared to a no-gRNA + DMSO control group (**Supplementary Figure 3**).

SunCLOUD9-mediated *SNAP91* loop formation in immature iGLUTs (DIV14) increased the ratio of neurites per neuron (*p* = 0.021, *r* = 0.22), and may have led to an increase in the number of branches formed per neurite (*p* = 0.074, *r* = 0.40) (**Figure 4E**). At a more mature timepoint (DIV21), there was a decrease in the intensity of SYN1+ synaptic puncta in those iGLUTs with the engineered loop (*p* = 0.024, *r* = 0.21) with suggestive evidence of an increase in the intensity of cell body puncta (*p* = 0.061, *r* = 0.17). These findings indicate that loop formation may have led to alteration of neurite branching patterns in immature neurons and a redistribution of synaptic puncta in iGLUT neurons with the *SNAP91* loop.

Consistent with these findings, spatial engineering of the *SNAP91* loop in iGLUT neurons led to marked differences in the development of mature firing properties. In the control condition, we observed the expected increase in population-wide weighted mean firing rate (WMFR) as neurons matured *in vitro*; furthermore, the duration of network bursting increased over time, with a concordant decrease in the frequency of such bursts (i.e., longer, less frequent network bursts as neurons matured). These changes were disrupted by creation of the *SNAP91* loop in iGLUT neurons, with a blunted rate of WMFR increase over time (*β* = -0.13 (-0.24 - -0.01), *p* = 0.032) and more frequent (*β* = 0.86 (0.28 – 1.45), *p* = 0.004) but shorter (*β* = -0.71 (-1.11 - -0.31), *p* = 0.001) network bursts **(Figure 4I, J)**. Furthermore, the network inter-spike-interval coefficient of variation (“Network ISI CoV”), a measure of spike variability within network bursts, decreased as the neurons matured in both groups; however, Network ISI CoV remained persistently elevated with induction of the *SNAP91* loop (*β* = 0.06 (0.02 – 0.09), *p* = 0.001). Finally, loop formation slowed the rate of maturation-dependent development of network synchrony in iGLUT neurons (*β* = -0.21 (-0.37 - -0.05), *p* = 0.010) **(Figure 4I, J)**. Taken together, these findings demonstrate a functional role for long-range, neuronal subtype-specific loop in regulating *SNAP91* gene expression and neuronal phenotypes.

### Demonstration of enhancer activity at a locus targeting a potential novel risk gene, BHLHE22

Towards functionally validating a loop involving a novel distal gene target of a schizophrenia risk locus, we identified an iGABA-specific loop involving the promoter region of *BHLHE22* and a distal non-coding region overlapping a risk locus 980kb away. Because *BHLHE22* is expressed at substantially higher levels in iGABA neurons than in the other two subtypes, this was a likely promoter-enhancer loop. The successful creation of the loop in DIV14 SunCLOUD9-iGLUTs was supported by an approximate three-fold increase in *BHLHE22* expression (*p* = 0.000054, *r* = 0.78).

At DIV14, *BHLHE22* SunCLOUD9-iGLUTs showed an increase in the average number of neurites per neuron (*p* = 0.054, *r* = 0.20) but a decrease in average neurite length (*p* = 0.0010, *r* = 0.34) and number of branch points (*p* = 0.00013, *r* = 0.39). The percentage of electrodes with bursting activity was consistently decreased *BHLHE22* SunCLOUD9-iGLUTs (*β* = -0.31 (-0.61 – 0.00), *p* = 0.049). Moreover, there was a significant interaction between DIV and treatment condition, such that *BHLHE22* SunCLOUD9-iGLUTs developed increased synchrony at a faster rate (*β* = 0.03 (0.00 – 0.05), *p* = 0.022). *BHLHE22* loop formation by SunCLOUD9 increased *BHLHE22* expression and impacted neuronal morphology and activity in iGLUTs.

## DISCUSSION

Comprehensive mapping of 3D chromatin structures in isogenic iDOPA, iGABA, and iGLUT neurons revealed the changing landscape of chromatin dynamics across neurodevelopment. Chromatin compartments and loops through which schizophrenia risk loci might alter neurodevelopmentally regulated gene expression programs in cell-type-specific manners are catalogued, highlighting a relatively new 3DG structure, architectural stripes, comprised of regions of brain-specific super-enhancer enrichments that connect schizophrenia risk loci to distal neurodevelopmentally regulated genes. Overall, we identified 167 neuron-specific stripes and 2920 loops specific to neuronal subtypes compared to either hiPSC or the other subtypes and predicted to connect schizophrenia risk variants and target genes across neurotransmitter systems. We discovered 263 risk genes contacted by “Risk Gene Connect Loops” that could potentially confer additional long-range regulation, and 2201 “Risk Locus Connect Genes”, of which approximately 1882 are potentially novel risk genes for schizophrenia. Unexpectedly, linear and 3D spatially proximal risk genes are enriched for distinct biological processes, neurotransmission and cell-cell adhesion, respectively. Finally, a novel chemical-induced proximity CRISPR system demonstrated the first-ever experimental validations of chromatin reorganization in human neurons: revealing the causal role of iGABA-specific loops targeting the proximal schizophrenia risk gene *SNAP91,* as well as a risk-locus-anchored region connected to a predicted distal schizophrenia risk gene *BHLHE22*.

3D genome structural mapping of chromatin^8, 12–18^, together with the activity-by-contact model^57, 58^, yield gene-enhancer maps that predict distal genes regulated through chromatin contacts. Although massively parallel reporter assays^59, 60^ and CRISPR perturbations can functionally verify promoter and enhancer activities at non-contiguous sequences^8, 9, 58, 61, 62^ and even resolve target genes (e.g. CRISPRi-FlowFish^58^), the role of spatial proximity on gene expression and neuronal function has not previously been definitively demonstrated. Here, we applied a SunCLOUD9-based approach^56^ in order to experimentally create chromatin loops linking regulatory regions anchored in risk loci to distal target genes. After confirming loop-formation and the resultant changes in target gene expression, we demonstrated the phenotypic consequences of manipulating chromatin loops on neuronal morphology and activity.

The neuronal phenotypes resulting from loop engineering were consistent with the biological roles of both genes studied. Rodent studies suggest a role for *SNAP91* in presynaptic function^51–54, 63^, and reduced expression of *SNAP91* in human neurons decreased synaptic activity^32^; here, a loop-based manipulation that lowered *SNAP91* expression likewise blunted the maturation-dependent increase in neuronal activity. *BHLHE22* is a transcription factor that regulates the developmental trajectory of neuronal projections in animal models^64, 65^; here, a loop-mediated increase in *BHLHE22* expression had potent effects on neurite length and branching patterns. To our knowledge, this represents the first successful application of chromatin loop engineering to post-mitotic neurons, demonstrating that the functional impact of the myriad 3DG structures associated with neurodevelopment and disease can now be empirically evaluated. Furthermore, our identification of a neuronal subtype-specific loop involving a schizophrenia-anchor enhancer region targeting *BHLHE22* through long-range interaction suggests that *BHLHE22* may represent a novel risk gene that would not have been identified without 3D genome techniques.

Translating SunCLOUD9 to a higher-throughput assay capable of validating hundreds of loops across multiple cell types is urgently needed, towards broadly exploring the role of 3D genome regulation. Leveraging pooled CRISPR-based screening approaches^66^ with high-throughput multi-omics single-cell readouts of chromatin structure^67^, open chromatin^68^, and gene-expression^69^ would extend this work and enable truly comprehensive screening of the developmentally-regulated, cell-type-specific and context-dependent target genes impacted by psychiatric risk loci.

Overall, the functions of proximity-based contacts between non-coding schizophrenia risk loci and distally targeted genes were probed, uncovering the importance of chromatin architectural stripes in cell-type-specific transcriptional regulation. Our neuronal subtype-specific 3D genome maps are accessible through the PsychENCODE Knowledge Portal (https://synapse.org) and further expand the number of Hi-C datasets from human brain^8, 9, 11, 17, 47, 70, 71^. Cell type–specific 3D genome analyses across the major neurotransmitter classes implicated in schizophrenia could improve the accuracy of diagnosis^72^ and the targets of distal risk genes may represent novel therapeutic targets^73^.

## AUTHOR CONTRIBUTIONS

SKP, WL, SA, and KJB conceived of the study. SKP, COS, SK, SG, RE, SH, PJMD and PA conducted experiments. SKP and COS prepared RNA-seq and Hi-C libraries. WQC and KCW designed the CRISPR-based loop editing tools, SKP and WL conducted computational and bioinformatic analyses. SKP wrote the paper with WL, KJB, and SA. All authors reviewed the manuscript and approved of it in its final form.

The authors thank Marina Iskhakova and Dana Infante for technical assistance

## DATA AND CODE AVAILABILITY

All source donor hiPSCs have been deposited at the Rutgers University Cell and DNA Repository (study 160; http://www.nimhstemcells.org/).

The source data described in this manuscript are available via the PsychENCODE Knowledge Portal (https://psychencode.synapse.org/). The PsychENCODE Knowledge Portal is a platform for accessing data, analyses, and tools generated through grants funded by the National Institute of Mental Health (NIMH) PsychENCODE program. Data is available for general research use according to the following requirements for data access and data attribution: (https://psychencode.synapse.org/DataAccess). For access to content described in this manuscript see: https://doi.org/xxxxxxx.

## METHODS

### Cell Culture

#### Human-Induced Pluripotent Stem Cell Culture

The hiPSC lines were derived via *OKSM* reprogramming of dermal fibroblasts with Sendai viral vectors and are from a previously established cohort^74^. hiPSCs were maintained in StemFlex media (Gibco, #A3349401) on matrigel-coated plates (Corning, #354230) and passaged with 0.5mM EDTA (Life Technologies, #15575-020) every four to seven days for a maximum of 10 passages. Monthly testing with the Lonza MycoAlert Mycoplasma Detection Kit (Lonza, #LT07-218) was used to confirm that all cultures were free of mycoplasma contamination.

#### Lentivirus Production

Third-generation lentiviruses *pUBIQ-rtTA* (Addgene #20342)*, tetO-ASCL1-LMX1B-NURR1-PuroR* (Addgene #182298), *tetO-ASCL1-PuroR* (Addgene #97329)*, tetO-DLX2-HygroR* (Addgene #97330), and *tetO-NGN2-eGFP-PuroR* (Addgene #79823) were generated using polyethylenimine (PEI, Polysciences, #23966-2)-mediated transfection of human embryonic kidney 293T (HEK293T) cells using existing protocols^75^. The plasmids and lentiviruses for the three SunCLOUD9 vectors were produced by VectorBuilder.

#### Production of Induced Dopaminergic (iDOPA), GABAergic (iGABA), and Glutamatergic (iGLUT) Neurons

All three neuronal subtypes were generated as described previously^25^. Induction relies on transient overexpression of lineage-promoting transcription factors combined with stringent chemical selection. iDOPAs were induced with *ASCL1*, *LMX1b*, and *NR4A2* ^25^ (also known as “Nurr1”); iGABAs with *ASCL1* and *DLX2* ^26, 27^; and iGLUTs with *NGN2*^33^. Briefly, hiPSCs were dissociated to single cells with Accutase Cell Detachment Solution (Innovative Cell Technologies, #AT104), quenched with DMEM (Gibco, #11965092), and centrifuged at RT at 800g for five minutes. Cell pellets were gently resuspended in StemFlex (Gibco, #A334901) supplemented with 10µM ROCK Inhibitor (StemCell Technologies, #72307). Equivalent titers of doxycycline-inducible lentivirus vectors encoding *tetO-ALN-PuroR* and *pUBIQ*-*rtTA* (iDOPA), *tetO-ASCL1-PuroR* and *tetO-DLX2-HygroR (iGABA),* or *tetO-NGN2-eGFP-PuroR (iGLUT)* together with *pUBIQ-rtTA,* were added to the suspension, mixed gently by inversion, dispensed onto Matrigel-coated plates, and incubated overnight at 37°C. The next day, media was changed to Induction Media supplemented with 1.0µg/mL doxycycline (DIV1), with antibiotic selections appropriate to the lentiviral vectors used (1.0µg/mL puromycin and 250µg/mL hygromycin (Thermo, #10687010) added the next day (DIV2) and continued for four days. 4µM Ara-C was included for four to six days after the initiation of antibiotic selection, and immature neurons were replated onto their final maturation plate by DIV7. For Hi-C, iDOPAs were matured until DIV35, iGABAs until DIV42, and iGLUTs until DIV21; timepoints reflected functional electrophysiologic maturation^25, 27, 33^.

#### Generation of hiPSC Lines Constitutively Expressing SunCLOUD9 Effectors

hiPSCs were transduced in suspension overnight with equal amounts of the three SunCLOUD9^56^ lentiviruses. Approximately 24 hours later, the media was replaced with StemFlex, and tripartite selection with 1.0 ng/mL puromycin, 0.25 µg/mL hygromycin, and 0.5 µg/mL neomycin began 24 hours afterwards and continued for 14 days, including at least one passage of hiPSCs in media with all three selection factors. Culturing donor- and passage-matched WT hiPSCs in the tripartite e selection media resulted in the complete elimination of all cells within five days, while numerous SunCLOUD9-transduced hiPSCs colonies survived and expanded. SunCLOUD9 hiPSCs were validated using a combination of rt-qPCR and immunocytochemical (ICC) techniques, as shown in **Supplementary Figure 1A-G**.

#### Generation of U6-gRNA-Expressing PCR Amplicons

Species-specific (*S. pyogenes* versus *S. aureus*) gRNAs targeting the two bins of the desired loop were designed in Benchling. The top three scoring sequences in each bin were selected for a total of six gRNAs per loop. The overlapping PCR primer approach was used as described in Ran et al., 2013^76^ to generate ∼350bp PCR amplicons expressing the desired gRNA sequence under the constitutively active U6 promoter (**Supplementary Figure 1H**). PCR amplicons were generated using the Agilent Herculase II Fusion Polymerase with dNTPs Combo Kit (Agilent, #600677). In brief, 50 µL reactions were set up containing 10 μM of a universal forward primer overlapping the first 20 base pairs of the U6 promoter, 10 μM of an amplicon-specific reverse primer containing the species-specific tracrRNA and the ∼20bp gRNA sequence, 10 µM a U6-promomter-containing plasmid template, Herculase Fusion Polymerase, PCR buffer, dNTPs (25mM each), and water. For each of the six reactions (one for each gRNA amplicon), a no-template water negative control was used to test for the production of primer-dimer artifacts. The reaction underwent 35 cycles, and the resulting products were purified using the QIAquick PCR Purification Kit (Qiagen, #28104) per the manufacturer’s instructions. Finally, the purified products were run for 30 minutes on a 2% agarose gel at 200V to ensure proper amplicon size around 350bps, the absence of primer artifacts in the no-template control, and the removal of the large (>10kb) plasmid template by the purification kit (**Supplementary Figure 1I**).

#### Engineering the iGABA-Specific Loops in SunCLOUD9-iGLUT Neurons

To evaluate the potential functional effects of iGABA-specific loops targeting the schizophrenia risk gene *SNAP91* or *BHLHE22*, the loop was created in iGLUT neurons and explored the impact on gene expression and relevant neuronal phenotypes. SunCLOUD9-hiPSCs were used to generate iGLUT neurons using the technique described above. On DIV1, 400ng of each of the six gRNA PCR amplicons was transfected per 10cm dish using the Lipofectamine 3000 Transfection Reagent per the manufacturer’s instructions (Thermo Fisher Scientific, #L3000001). 500μM ABA or a volumetric equivalent of DMSO was added to the media starting on DIV2 and continued until DIV14, with full media replacements daily (**Supplementary Figure 1J**).

### Neuronal Phenotypic Analyses

#### RNA Extraction and Reverse-Transcription qPCR

Cells were washed twice with PBS and lysed with TRIzol Reagent (Thermo, #15596026). The Direct-zol RNA miniprep kit with in-column DNAse treatment (Zymo Research, #R2051) was used to isolate and purify RNA. For rt-qPCR, 50ng technical quadruplets were loaded into a 384-well plate and quantified using the *Power* SYBR Green RNA-to-C_t_ *1-Step* Kit (Thermo, #4389986). Relative mRNA abundance levels were determined using the ΔΔ-C_t_ method.

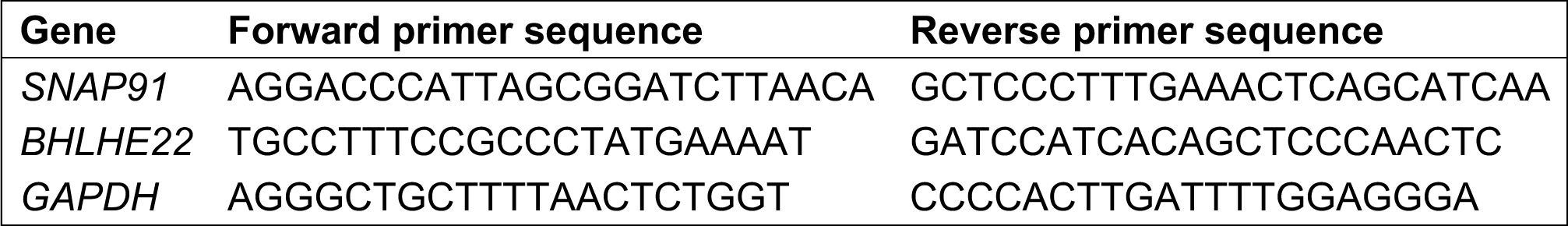

#### Immunocytochemistry

In brief, cells were plated onto glass coverslips in a 24-well plate until the desired timepoint. Cells were washed twice with PBS and fixed with 4% formaldehyde in PBS (Electron Microscopy Sciences, #15170), followed by three PBS washes. Cells were blocked for one hour at room temperature with 5% donkey serum (Jackson, #017-000-121) and 0.1% Triton X-100 (Sigma, #T8787), washed three times, and incubated overnight at 4 °C with primary antibodies in 5% donkey serum and 0.1% Tween-20 (Boston BioProducts, #IBB-181X) in PBS. Cells were washed twice, incubated with the appropriate secondary antibodies in PBS in a dark room at 4 °C, washed three additional times, incubated in 0.5 µg/mL DAPI (Sigma, #D9542) for 5 minutes at RT, and washed two more times in PBS.

| Antibody | Species | Supplier | Product number | Dilution |
| --- | --- | --- | --- | --- |
| MAP2 | Ck | Abcam | ab5392 | 1:500 |
| SYNAPSIN | Ms | Synaptic Systems | 106 011 | 1:500 |
| TUJ1 | Ms | BioLegend | MMS-435P | 1:1000 |
| Alexa 568 anti-Mouse | Ms | Life technologies | A10037 | 1:500 |
| Alexa 647 anti-Chicken | Ck | Life technologies | A10042 | 1:500 |

#### Multi-Electrode Array (MEA)

DIV7 iGLUT neurons were plated onto 48-well matrigel-coated MEA plates (Axion Biosystems, M768-tMEA-48W) containing commercially available human astrocytes (HA; Sciencell, #1800) and maintained them in Neuron Media supplemented with 2% heat-inactivated fetal bovine serum (FBS, Gibco, #16140071) throughout the experiment. 2µM Ara-C was included in the media for the first week to arrest HA division. Half media changes were performed twice a week, one day before MEA measurement; MEA measurements began on DIV14 at the earliest and took place 2-3 times per week on a Maestro Multi-Electron Array system (Axion Biosystems). Prior to starting the recording, the plate was equilibrated in the machine for five minutes, and data were collected for a total of 10 minutes, with spontaneous neural real-time configuration at a threshold of 5.5. After the final timepoint, all recordings for a given experiment were batch-processed. For each electrophysiologic feature of interest, we fit log-transformed linear mixed effects models, with random slopes assigned for each well.

#### High-Content Imaging

hiPSCs were transduced with the lentiviruses needed to generate the desired neuronal subtype using the methods described above. After ∼7 days of selection with the appropriate antibiotics and Ara-C, the cells were split and replated on 96-well plates. For neurite-tracing, cells (∼25,000 per well) were plated on 80 µg/mL matrigel and imaged on DIV14 or DIV28. At the appropriate timepoint, we performed ICC staining of the neurons using the protocol detailed above. Cells were stained for TUBIII and MAP2 for neurite-tracing and TUBIII and SYN1 for synaptic imaging. Immunostained plates were imaged with the *CX7 High Content Analysis Platform* (Thermo Fisher). For neurite-tracing, at least four wells in each condition, with at least 9 fields of view in each well, were imaged at 20X resolution. Wilcoxon-Mann-Whitney tests were used to assess potential group differences using the *wilcoxon_test()* function from the R packages *RStatix* and *coin*. A *p*-value threshold of < 0.05 was used to determine statistical significance.

### 3D Genome Assays

#### Chromosome Conformation Capture (3C)

At least 10 million neurons were harvested per sample for fixation in 1.5% formaldehyde solution, followed by a quench in 0.125 M glycine, pelleting by centrifugation, and lysis using a Dounce Homogenizer in 3C Lysis Buffer. Chromatin was pelleted, washed it twice with 1X NEB Buffer 2.1 (New England BioLabs, #B7202), diluted it in buffer, followed by removal of non-crosslinked proteins via incubation in SDS for 10 minutes at 65 °C. Restriction enzyme incubation took place overnight at 37 °C, after which the enzyme was inactivated via incubation in SDS for 30 minutes at 65 °C. Samples were diluted in freshly made 3C Ligation Mix, incubated them for 2 hours at 16 °C, and reversed crosslinks by incubating overnight with 50µL of 10mg/mL Proteinase K solution (Thermo Fisher, #25530031) at 65 °C, followed by an additional incubation after adding 50µL more of the Proteinase K solution. DNA was isolated via phenol:chloroform extraction and precipitated in 1X TE Buffer. Finally, residual RNA contamination was removed by incubating the samples in 10µL of 1mg/mL recombinant RNase (Ambion, #AM2269). Sample aliquots of 100µL were kept at -80 °C.

3C templates (at least two experimental replicates) were amplified using the selected primer pairs and run on a 2% agarose gel in triplicate. Semi-quantification of band intensity was performed with imageJ and normalized the values to the intensity of bands from a DNA loading control (LC) produced via amplification of 3C templates with the PC1/PC2 primer pair. The sample LC-normalized band intensity was divided by the LC-normalized band intensity produced with a no T4 ligase negative control condition from the corresponding sample to generate adjusted interaction frequency values, which were combined by sample identity for all selected primer pairs amplifying the same or adjacent restriction fragments (including the two experimental replicate 3C template libraries and the DNA electrophoresis replicates). Statistical analysis was conducted with Wilcoxon-Mann-Whitney tests using the *wilcoxon_test()* with the *RStatix* package in R, and effect sizes were determined via the *wilcoxon_effsize()* function of the *coin* package in R.

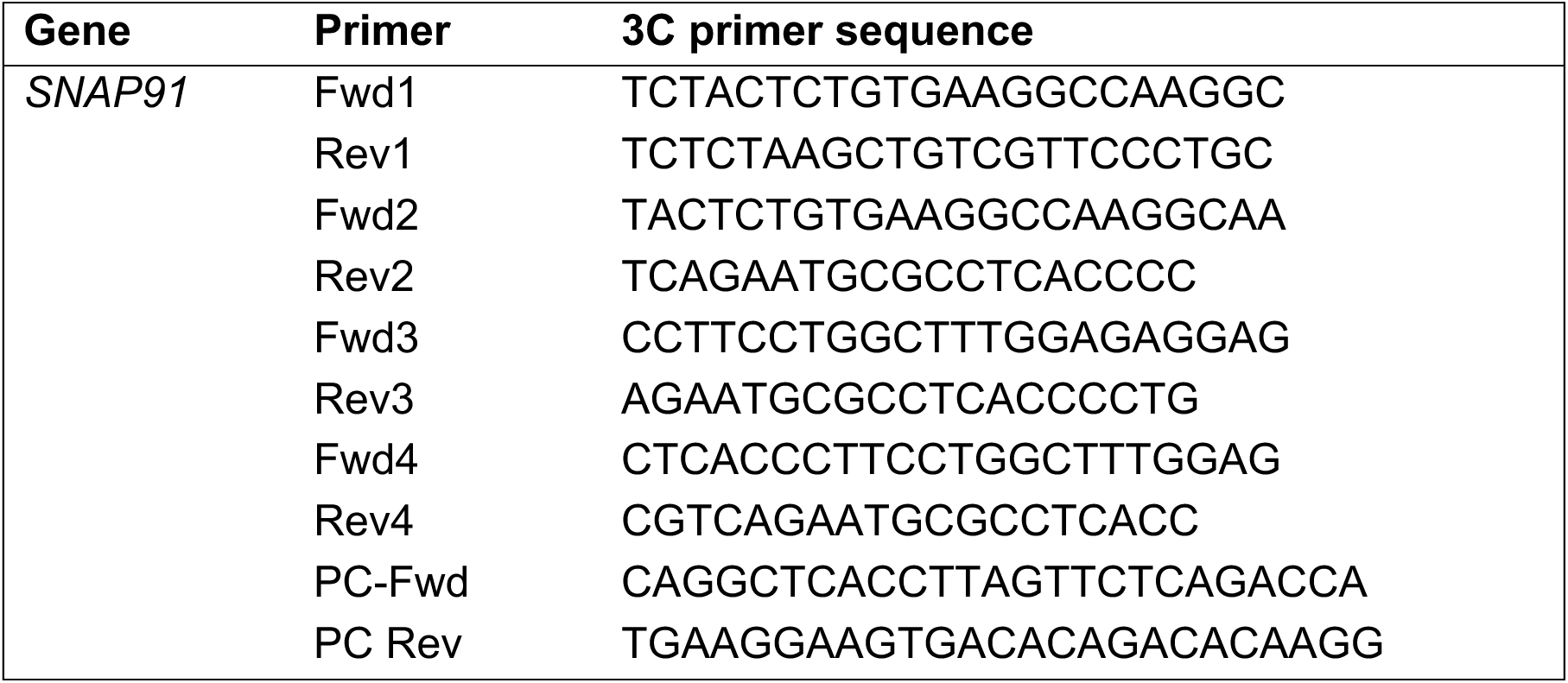

#### Genome-Wide Chromosome Conformation Capture (Hi-C) Sample-Processing

Samples were processed to generate Hi-C libraries according to the protocol for the *Arima Hi-C Kit User Guide for Mammalian Cell Lines* (Arima Genomics, #A510008). Cells were dissociated by incubating in Accutase (Stemcell Technologies, #07920) for ∼10 minutes at 37 °C. The cell suspension was quenched with DMEM (Gibco, 11965) at a volumetric ratio of 1:3 and pelleted by centrifugation at room temperature for 5 minutes at 1,000 x g. Cells were resuspended in 1mL PBS and counted via Trypan Blue exclusion on a Countess machine. Aliquots of at least 2 million cells were transferred to new 15mL Falcon tubes and PBS was added to 5mL to wash. Cells were then pelleted and resuspended 2% formaldehyde in PBS. Samples were incubated at room temperature for 10 minutes while rotating. Stop Solution 1 was added to quench formaldehyde for 5 minutes at room temperature while rotating. Fixed cells were once again pelleted and washed by resuspension PBS, pelleted, and were lysed via resuspension Lysis Solution for 15 minutes at 4 °C. Chromatin was digested with the kit’s proprietary restriction enzyme cocktail. 5’ DNA overhands were filled in and labelled with biotinylated nucleotides. Intramolecular DNA-ligation and crosslink-reversal were conducted per the manufacturer’s instructions. We used AMPure XP Beads (Beckman Coulter, #10136224) to pull down and purify ligated DNA fragments, using two washes with 80% ethanol, followed by elution using the kit’s Elution Buffer.

#### Generation of Hi-C Libraries for Sequencing

Eluted libraries were brought to 100uL with Elution Buffer and fragmented using a Covaris sonicator with a target mean fragment size of 400bp. Size selection (200-600bp) was performed with AMPure XP Beads, and samples were washed twice with 80% ethanol on a magnetic rack and eluted in Elution Buffer. Samples were enriched for biotinylated fragments using the kit’s Enrichment Beads followed by washes in kit Wash Buffer. End repair was conducted per the manufacturer’s instructions. Adapters were ligated using the Swift Biosciences Accel-NGS 2S Plus DNA Indexing Kit (Swift Biosciences, #21024). Following adapter-ligation, washes, and elution, the libraries were amplified using the KAPA Library Amplification Kit (Roche, KK2702) with 12 PCR cycles for amplification. Finally, samples were purified with AMPure XP Beads and eluted in Elution Buffer. Library concentration was assessed using both Qubit fluorometric quantification and the qPCR-based KAPA Library Quantification Kit (Roche, KK4824). Appropriate fragment size distributions were confirmed by running each sample on an Agilent Bioanalyzer using the DNA High-Sensitivity Kit (Agilent, #5067-4626).

#### Sequencing of Hi-C Libraries

Hi-C libraries were pooled into four libraries per group (two groups total) and subjected to 75bp PE sequencing on a NovaSeq 6000 system (Illumina), with each group occupying a whole lane of an S4 sequencing kit, leading to a predicted sequencing depth of 500 million reads per library.

### Computational Analyses of Sequenced Hi-C Libraries

#### Primary processing

Initial processing of the raw 2×125bp read pair FASTQ files was performed using the HiC-Pro v2.11.1 analysis pipeline^77^. In brief, HiC-Pro performs four major tasks: aligning short reads, filtering for valid pairs, binning, and normalizing contact matrices. HiC-Pro implements the truncation-based alignment strategy using Bowtie v2.2.3^78^, mapping full reads end-to-end or the 5’ portion of reads preceding an appropriate ligation site that results from digestion with restriction enzyme cocktail used in the *Arima Hi-C Kit User Guide for Mammalian Cell Lines*. Invalid interactions such as same-strand, dangling-end, self-cycle, and single-end pairs are not retained. Binning was performed in non-overlapping, adjacent windows across the genome and resulting contact matrices were normalized using iterative correction and eigenvector decomposition (ICE) as previously described^79^. HiC-Pro outputs were converted to *cool* files^80^ using *hicConvertFormat* from the HiCExplorer^81^. Matrices were balanced using *‘cooler balance’*, masking ENCODE blacklisted regions^82^. To create high coverage *cool* files for each cell type, iDOPA, iGABA, iGLUT, and hiPSCs, data from individual donors were merged and down-sampled to match the sample with the lowest number of intrachromosomal contacts using ‘*cooltools random-sample’*^83^.

#### Compartments and topologically associated domains (TADs)

To identify boundaries of low-resolution A/B compartments associated with open and closed chromatin, respectively, ‘*cooltools call-compartments’* was used to perform eigenvector decomposition as implemented in the 4D Nucleome (4DN) analysis workflow docker image^84^. Similarly, TADs were identified using ‘*cooltools diamond-insulation’*^85^.

The developmentally regulated A/B compartment map was super-imposed with the most recent genome wide association study (GWAS) map for schizophrenia^3^, comprised of 291 common risk loci and 1111 PsychENCODE schizophrenia risk genes (http://resource.psychencode.org/Datasets/Integrative/INT-18_SCZ_Risk_Gene_List.csv).

#### Loops and stripes

To identify loop interaction hotspots, we employed the supervised machine-learning loop-caller, Peakachu^46^, using the appropriate CTCF model for the given intrachromosomal reads in each sample with a 0.9 probability threshold cut-off. Stripes were detected using StriPENN^43^ with default parameters, ‘*stripenn compute*’. Stripes were initially called for each high coverage cell-type *cool* file. Contiguous stripes were then consolidated into a single, nonoverlapping set and scored for each of the high-coverage cell-type *coolers* with ‘*stripenn score’*.

#### Significant interactions with specific bin(s)

To identify significantly enriched interactions involving a bin or a set of bins comprising loci of interest with another bin, we employed a previously used method^13^, adapted from a procedure proposed by Won, H. et al^8^. In brief, the expected interaction counts for each interaction distance was estimated by calculating the mean of all intrachromosomal bin-bin interactions of the same separation distance throughout the raw intrachromosomal contact matrix using the R package, HiTC^86^, to facilitate manipulation of our HiC-Pro-produced raw contact matrices and estimation of the expected counts at various interaction distances. The probability of observing an interaction between a bin-of-interest and another bin was then defined as the expected interaction between those two bins divided by the sum of all expected interactions between the bin-of-interest and all other intrachromosomal bins. A *p*-value was then calculated as binomial probability of observing the number of interaction counts or more between the bin-of-interest and some other bin where the number of successes was defined as the observed interaction count, the number of tries as the total number of observed interactions between the bin-of-interest and all other intrachromosomal bins, and the success probability as the probability of observing the bin-bin interaction estimated from the expected mean interaction counts. The Benjamini-Hochberg method was used to control false discovery rate (FDR) for p-values determined for all interactions with a bin-of-interest (includes all bins 1Mb up and downstream in our tests).

## SUPPLEMENTARY FIGURES

**Supplementary Figure 1:**
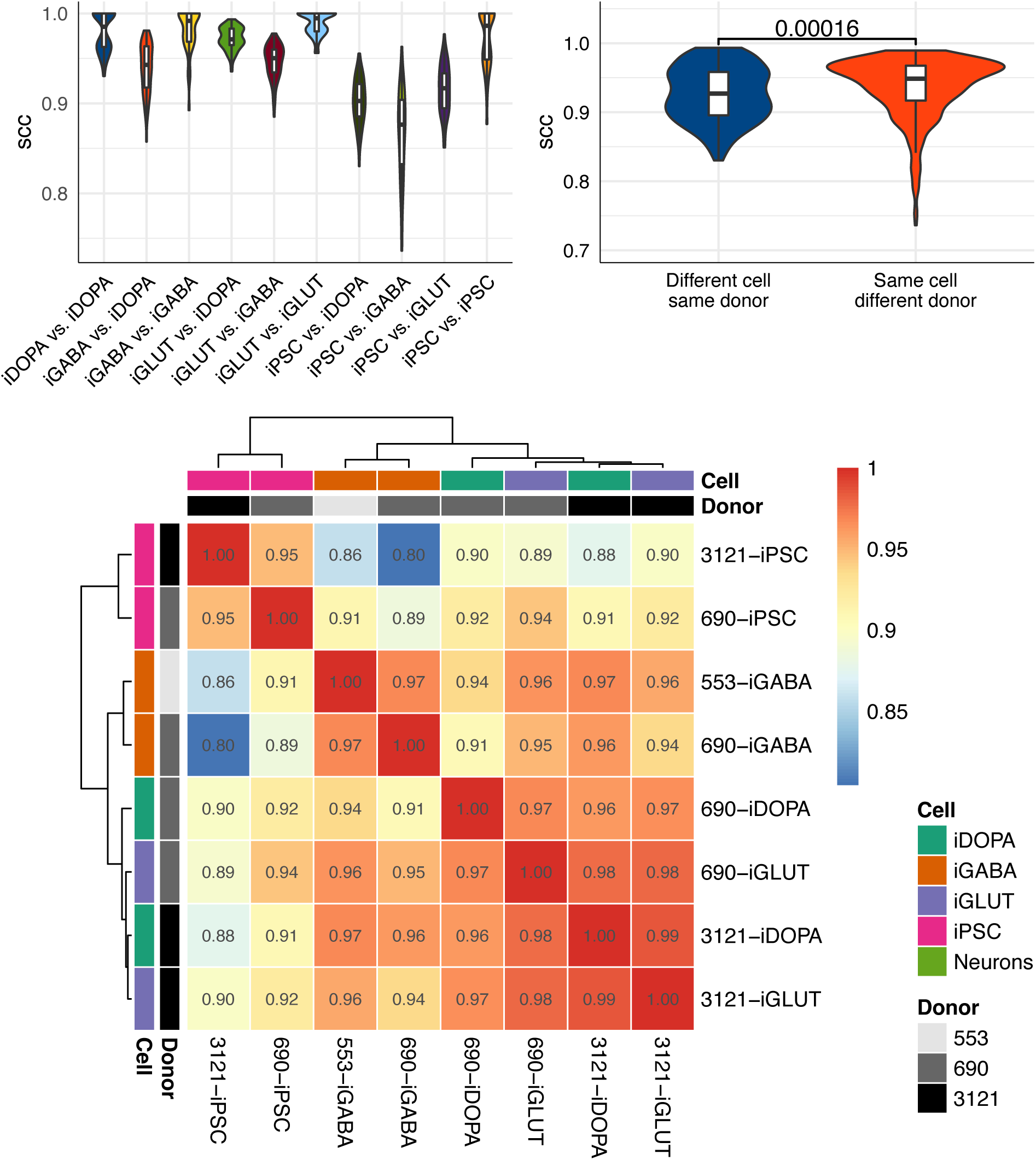
Hi-C-style correlation between donor contact maps. Similarity between samples, at the donor and cell type level, using stratum adjusted correlation coefficient (SCC) metrics. There was substantially higher SCC within cell types compared to within donors (*p* = 1.6×10^-4^), as well as between same cell-types compared to different cell types.

**Supplementary Figure 2:**
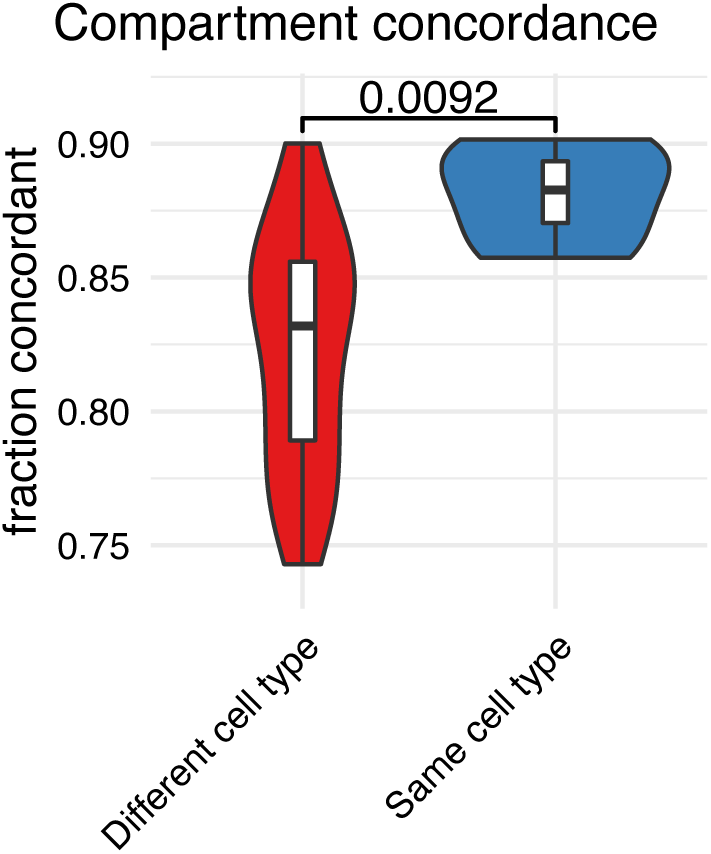
Agreement between donors on compartment calls. Chromosomal compartment architectures within a given cell type were highly reproducible across donors, the fraction of concordant compartment calls between same cell types was significantly greater then between different cell types (*p* = 9.2×10-3).

**Supplementary Figure 3:**
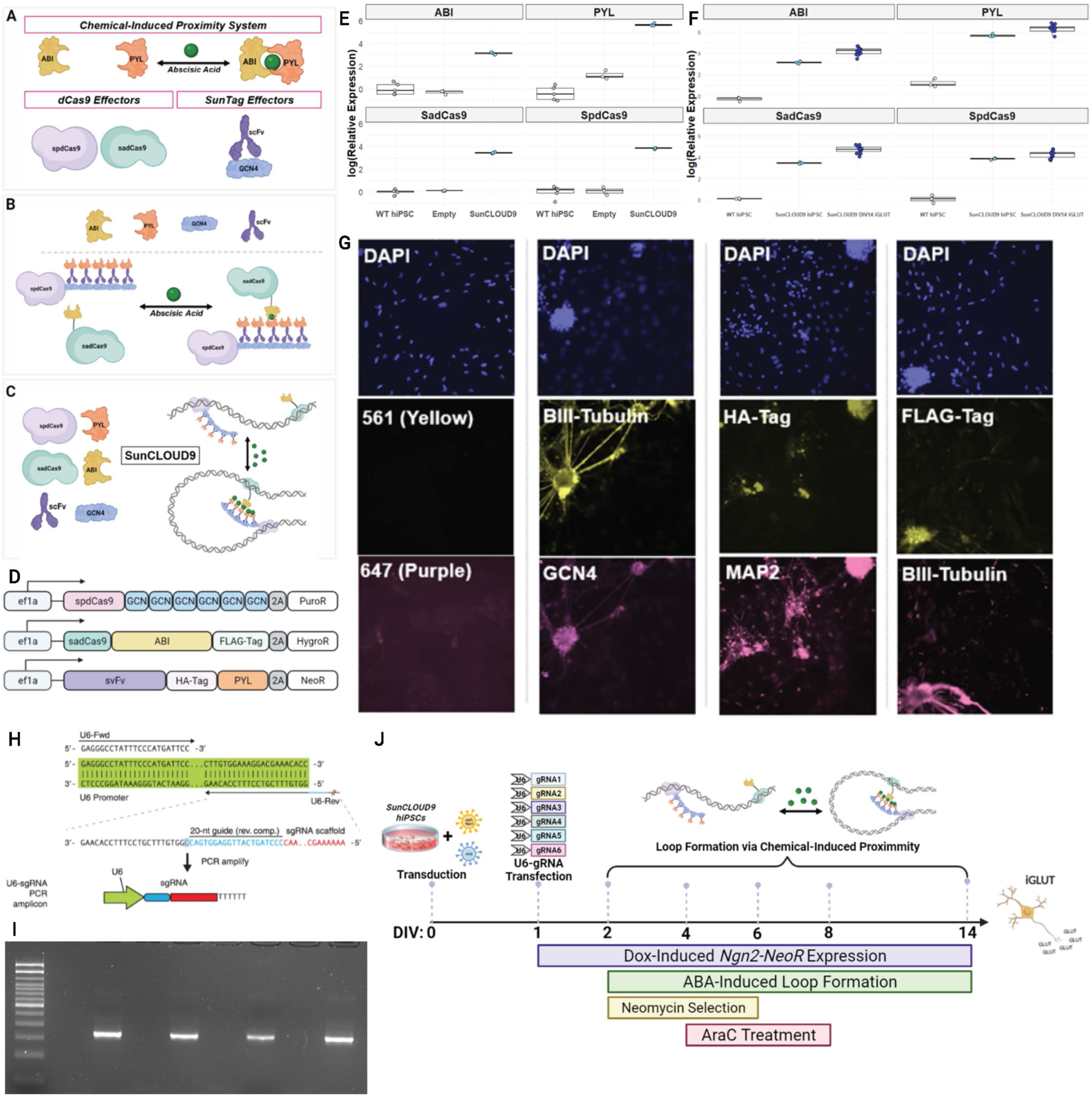
Technical Development of the SunCLOUD9 Method for Chromatin Engineering. **(A)** Schematic illustration of the key effectors of the SunCLOUD9 system^55, 56^. Chemical-induced proximity is achieved through abscisic acid (ABA) and the plant dimerization domain ABI + PYL. **(B)** SunCLOUD9 fusion protein effectors after translation. **(C)** ABA-induced physical proximity of chromatin loci targeted with spdCas9-24XGCN4-scFv-PYL and sadCas9-ABI. Induced physical proximity of chromatin loci is chemically reversible upon removal of ABA. **(D)** Schematic of SunCLOUD9 lentivirus vectors. **(E)** SunCLOUD9-hiPSCs show mRNA expression of key effectors at several log-fold increases above background signal observed in WT hiPSCs and hiPSCs that received an empty transfection. **(F)** mRNA expression of SunCLOUD9 effectors in transgenic hiPSCs and DIV14 iGLUT neurons. **(G)** Immunocytochemical analysis of SunCLOUD9 transgene expression in DIV14 iGLUTs. The far-left column represents a secondary-only negative control. iGLUTs are co-cultured with human astrocytes lacking SunCLOUD9 transgenes, serving as an internal, negative biological control. **(H)** Schematic of PCR amplicon generation. **(I)** Representative agarose gel showing U6-gRNAs (350bp) targeting the *SNAP91* loop. Lanes alternative between reactions with and without the template to test for primer-dimer artifacts. **(J)** Timeline for SunCLOUD9-mediated engineering of chromatin loops in iGLUT neurons. hiPSCs stably expressing the SunCLOUD9 effectors are transduced with tetO-Ngn2-NeoR and rtTA lentiviruses. Induction of NGN2 and transfection of U6-gRNA PCR amplicons at DIV1. Addition of ABA for chemical-induced proximity at DIV2.

## SUPPLEMENTARY DATASETS

**SI Dataset 1: ‘S1. Hi-C QC.xlsx’**

A. Donor-level QC
B. Donor-merged QC (downsampled to 185 million *cis* contacts counting stats)

**SI Dataset 2: ‘S2. Compartments.xlsx’**

A. Donor-level calls
B. Donor-level eigenvector1
C. Merged calls
D. Merged eigenvector1
E. DEGs in switch comps
F. gProfiler for switch DEGs

**SI Dataset 3: ‘S3. Stripes.xlsx’**

A. Donor-level stripes
B. Differential stripes
C. DEGs in diff stripes
D. gProfiler on stripe DEGs
E. SEs (super-enhancers) in diff stripes

**SI Dataset 4: ‘S4. Loops.xlsx’**

A. Combined loop calls
B. Enriched GO:BP in neu loops (neuron subtype specific vs. other subtypes)
C. Simplified GO:BP in neu loops
D. Risk/Non-risk GO:BP
E. Simplified Risk/Non-risk simple GO:BP

**SI Dataset 5:’S5. expression.txt’, differential expression summary.**

